# Stratified neural networks in a time-to-event setting

**DOI:** 10.1101/2021.02.01.429169

**Authors:** Fabrizio Kuruc, Harald Binder, Moritz Hess

## Abstract

Deep neural networks are now frequently employed to predict survival conditional on omics-type biomarkers, e.g. by employing the partial likelihood of Cox proportional hazards model as loss function. Due to the generally limited number of observations in clinical studies, combining different data-sets has been proposed to improve learning of network parameters. However, if baseline hazards differ between the studies, the assumptions of Cox proportional hazards model are violated. Based on high dimensional transcriptome profiles from different tumor entities, we demonstrate how using a stratified partial likelihood as loss function allows for accounting for the different baseline hazards in a deep learning framework. Additionally, we compare the partial likelihood with the ranking loss, which is frequently employed as loss function in machine learning approaches due to its seemingly simplicity. Using RNA-seq data from the Cancer Genome Atlas (TCGA) we show that use of stratified loss functions leads to an overall better discriminatory power and lower prediction error compared to their non-stratified counterparts. We investigate which genes are identified to have the greatest marginal impact on prediction of survival when using different loss functions. We find that while similar genes are identified, in particular known prognostic genes receive higher importance from stratified loss functions. Taken together, pooling data from different sources for improved parameter learning of deep neural networks benefits largely from employing stratified loss functions that consider potentially varying baseline hazards. For easy application, we provide PyTorch code for stratified loss functions and an explanatory Jupyter notebook in a GitHub repository.

## General introduction

In biomedical applications, frequently time-to-event data is investigated. A common task is, e.g., to perform risk stratification of patients based on molecular profiles, such as retrieved by querying the transcriptome of tumor tissue [1] [2]. Today, multivariable approaches tailored to time to event data such as survival random forests [3], support vector machines (SVMs) [4] or approaches building on the concept of boosting [5] [6] are routinely applied for this task. However, recent advances in deep neural networks promise to uncover complex dependencies in the data and thereby might allow for better predictions of survival, e.g., conditional on gene expression profiles [7] [8] [9] [10].

Analyzing time-to-event data differs fundamentally from other data types. Right-censoring and potentially complex distributions of survival times makes use of fully parametric models difficult. Instead the Cox proportional hazards model [11] is frequently applied. Here, the instantaneous risk of observing an event at a given point in time is factorized into the baseline hazard function and a term with a linear predictor that indicates the proportional risk of observing a hazardous event based on the values of covariates, e.g., the expression of genes, for a given observation. Only the linear predictor is parameterized in Cox model, while the baseline hazard is not inferred. Correspondingly, a partial likelihood is used for parameter estimation.

Fitting neural networks based on the gradient of the partial log-likelihood function of the proportional hazards model allows to learn the potentially complex interplay of covariates in predicting the relative probability of a hazard conditional on values observed for the covariates. This approach has been employed in many studies, e.g. in [12], [10] or [7].

Still, biomedical data sets often comprise a rather small number of observations, mostly due to time-consuming clinical annotations or difficulties in getting e.g. tumor tissue. For improving the learning of deep neural networks, solutions such as proposed in [7] allow to jointly analyze multiple clinical data sets, e.g., originating from different tumor entities.

However, different tumor entities can differ substantially in their baseline hazard rate, which might introduce a bias, as the proportional hazards assumption, which is inherently related to factoring out the baseline hazard, is violated [13] [14]. This can adversely affect the prognostic relevance of a marker, especially with increased observation time [13].

To jointly analyze data sets differing in their baseline hazard, stratified proportional hazards models have been proposed (see [15] for details). Yet, this established approach from biostatistics surprisingly has not found uptake in the neural network community. Therefore, our aim is to evaluate the extent of potentially lost prediction performance due to this omission, and more generally to describe how stratification can be adequately implemented.

Specifically, we also investigate approaches based on the concordance index (C-index). This is a performance criterion that is frequently used for evaluating Cox proportional hazards models [16]. This criterion indicates, whether the ranks of observed hazard times are in concordance with the ranks of the risk scores computed for the subjects that experience a hazard at a given time. The C-index can be seen analogous to the area under the curve of a binary classification model [17]. A loss that directly maximizes the C-index is the ranking loss. Due to the seemingly straightforward objective, the ranking loss is frequently employed instead of the partial likelihood in some of the afore mentioned machine learning approaches, e.g. in [4] [6] [8]. Yet, the C-index only indicates the accuracy of classification, and not the calibration of the model [18]. Therefore, we not only propose and evaluate a stratified version of the C-index as a loss function, but consider performance of the C-index as a loss function per se.

For straightforward application of the methods, we provide the code for stratified loss functions, implemented in PyTorch, and an explanatory Jupyter Notebook in a GitHub repository.

## Artificial neural networks in a time-to-event setting

Artificial neural networks are capable of approximating even very complex functions [20]. This makes them ideally suited to solve a supervised learning task corresponding to learning some function *f* from given data (*x_i_, y_i_*), where *x_i_* describes the features and *y_i_* the corresponding label of a data point. Fully connected feed-forward networks are built of nodes arranged in *l* consecutively arranged hidden layers *h*, where each layer emerges from an affin-linear transformation and a non-linear activation function *φ*, component-wise applied on the preceding layer. A common choice for *φ* is the Rectified Linear Unit (ReLU). The first hidden layer operates on the input data. The activation of the nodes in each hidden layer (*j*) can be computed according to:

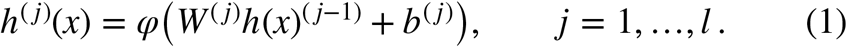

The parameters of the network are *W*and *b* and their shapes correspond to the number of nodes in each layer. The number of hidden layers indicates the depth. Given eq. 1, the network generates an output

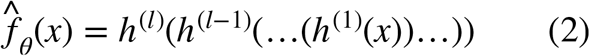

where *θ* = (*W*^(1)^, *b*^(1)^,…, *W*^(*l*)^, *b*^(*l*)^). This output is then used in a loss function together with the corresponding labels of the input data (see fig. 1 B). A natural choice is the negative log-likelihood function of the assumed model. The parameters are then iteratively approximated through back-propagation of errors [21] as returned by the loss function.

**Figure 1:**
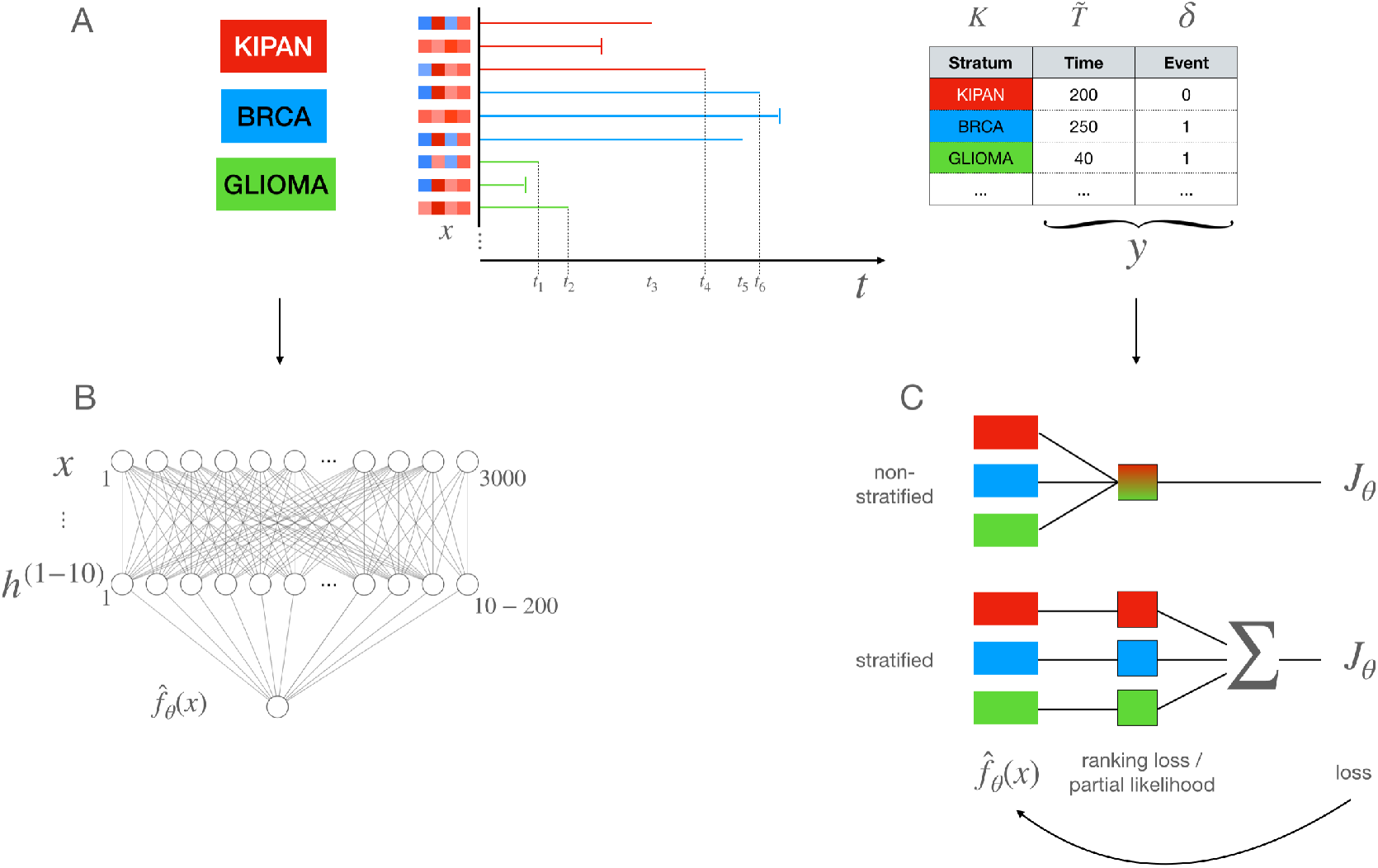
Predicting survival conditional on gene expression with deep neural networks trained with stratified and non-stratified versions of ranking loss and partial likelihood. (A) Gene expression data, extracted from tumors, as well as the time until a patient died (the event) are available from the TCGA. The aim is, to predict, if a patient survived until a given point in time. Event times 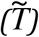
 and corresponding expression profiles (x) of four hypothetical genes (expression is shown standardized) are shown for nine hypothetical patients. End of follow-up without an experienced event is indicated by vertical bars (censored observations; δ = 0). Additionally the cancer subtype (the stratum; K) is provided. (B) Feed forward networks with a varying number of hidden layers (h) and nodes per hidden layer, are trained on the standardized expression data of the 3000 most variable genes. They learn the latent representation 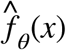
 of the input data, given the loss (J_θ_) computed by the functions mentioned in C. (C) We investigate non-stratified and stratified versions of commonly employed loss functions, specifically the ranking loss and the partial likelihood. In the stratified loss functions, the loss is first calculated within the different strata and then combined to adjust the weights of the neural network shown in B by back-propagation of errors.

In a time-to-event setting, the data consists of an individual feature vector *x* and, in the case of right censored data, of a composite label. The first part of the label is the observed event time 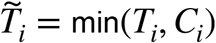, which can be either the occurrence time of the actual event or the censoring time (*C_i_*). The second part is a binary value indicating whether an event was observed at the time or not *δ¿ = I*(*T_i_* ≤ *C¿*) (where *I*(·) is the indicator function) (see fig. 1 A). A common model in this setting is Cox proportional hazards model [11]. It uses a semi-parametric approach and assumes a multiplicative relation for the conditional hazard of experiencing the event (*λ*(*t* | *x*)) between an unspecified baseline hazard function (*λ*θ(*t*)) and a parametric transformation of the individuals features (exp(*β′x*)):

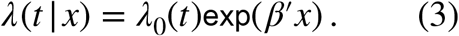

As the baseline hazard typically is left unspecified, there is no full likelihood for optimization. However, parameters can still be consistently approximated through a partial likelihood. Replacing the linear combination of parameters and features (exp(*β’x*)) in the negative log partial likelihood function through the output of a neural network, we retrieve the loss function for our survival data, which then can be used in the back-propagation of errors:

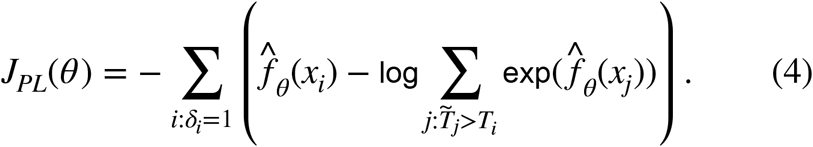

## Stratified loss functions for time to event data

### Stratified partial likelihood

The fundamental assumption of the proportional hazards model regards the baseline hazard function, which is assumed to be the same for every individual in the data set. Estimation will be biased if this assumption is violated, e.g., if distinct sub-groups are present due to different tumor types being jointly analyzed. To account for different risk groups in the data, stratification can be carried out along the strata that are assumed to violate the assumption. If the proportional hazards criterion is met within the identified groups, estimation can be carried out consistently again. Transferring this situation into a neural network time-to-event setting, the stratification procedure can be applied through the used loss function. The overall loss function will then just be the accumulation of calculated loss from each group (see fig. 1 C). Therefore the formulation of a stratified version of the partial likelihood loss is straightforward:

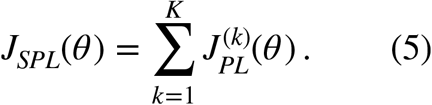

Note that the conventional partial likelihood is just a special case with *K* =1.

### Stratified ranking loss

In a time-to-event setting a ranking problem is concerned with the sorting of survival times as closely as possible to the actual true order. Ranking can be carried out through assigned risk scores, which in the case of the proportional hazards model is given through the linear combination of fitted parameters and covariates as:

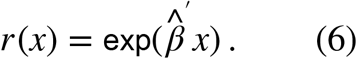

In order to maximize the correct ranking of survival times, a metric is needed to access the discriminatory power of the model. The C-index is a widely used metric for this purpose. It approximately reflects the probability that for a randomly selected pair of individuals, the person with a higher risk score has a shorter survival time [22]. Event times of censored individuals cannot be compared with larger survival times, as it is not clear who survived longer. Thus, the set of valid comparable individuals reduces to 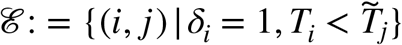
 and the C-index is given through:

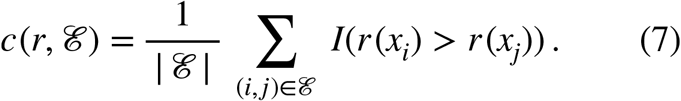

The maximization of this objective within a neural network setting requires to deal with the non-differentiable indicator function of the concordance index. In order to incorporate stratification into this setting the set of valid comparable individuals must be additionally calculated within each risk group. A stratified ranking loss function is therefore given by:

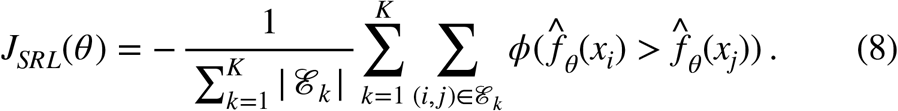

Again, the non-stratified version is just a special case of this function with *K* =1. Note that the linear combination of risk scores is replaced by a non-linear dependence through the replacement with the output of the neural network. Further, the indicator function is here replaced by *φ*, which must be a differentiable lower bound. By minimizing the whole objective function the negative of the concordance index is minimized and therefore the ranking metric maximized. An overview over applicable functions can be found in [17].

### Evaluation criteria

#### Stratified C-index

For the evaluation of the conventional concordance index in a proportional hazards setting, the linear combination of fitted values and features can be used, as the same baseline hazard is assumed for all individuals and the exponential function is a strictly monotone increasing function. A stratified setting allows for different baseline hazards within each risk group and therefore this approach cannot be taken. To adjust for this circumstance the set of valid comparable individuals has to be calculated within each subgroup, as this are the groups where we assume the same baseline hazard for all individuals, such that we can again compare risk scores through the yielded linear combination of features and parameters. Thus, a stratified version of the concordance index is given through:

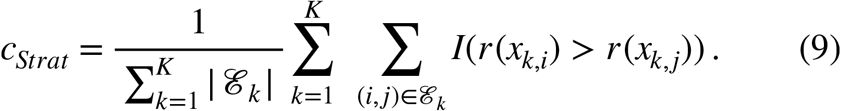

#### Stratified prediction error

The C-index explicitly targets the discriminatory power of a model. As this criterion is invariant to monotone transformations, it can not distinguish between well and badly calibrated models and suffers additionally from a bias induced by right censoring [23]. A perfectly discriminating model can be poorly calibrated and vice versa [18]. One alternative criterion suggested is the expected prediction error, which is also known as Brier score. It takes both, calibration and discrimination into account and therefore can be regarded as an overall performance measure, which can be seen in more detail through a decomposition of the expected Brier Score [24], [25], [19].

For a fitted neural network with stratified proportional hazard loss, the probability for an individual to survive until time *t* is estimated as:

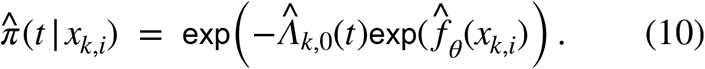

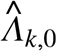
 is the Breslow estimator of the cumulative baseline hazard 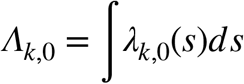
 of risk group k. This estimate is given through:

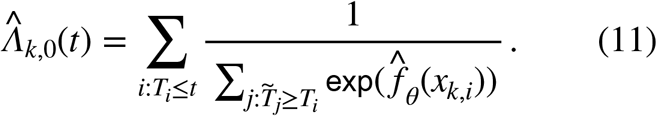

The true prediction error is given as the expected difference of the true status and the predicted survival probability of the individuals at time t according to:

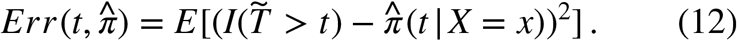

When empirical estimates of the prediction error are considered, inverse probability of censoring weights (IPCW) allow to take censoring into account for a consistent estimation [19], [25]. As we are considering the case with different risk groups present, the censoring distribution has to be estimated within in each subgroup. The prediction error for the validation data is therefore obtained as:

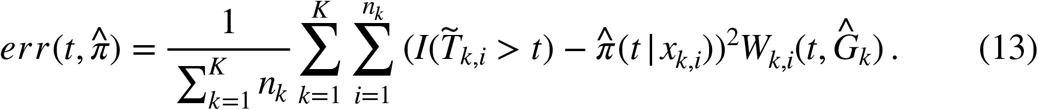

Note that the estimate is normalized through the total amount of individuals to obtain values between 0 and 1. The sum of the weights divided by the amount of total observed individuals sums up to 1. The censoring weights for each risk group are given through:

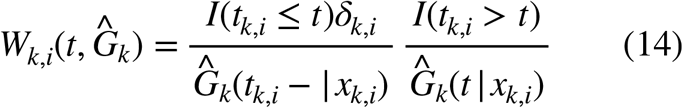

and depend on an consistent estimate 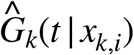 of the censoring distribution 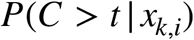, which in turn depends on the covariates 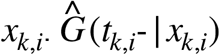 indicates the time just before *t* conditioned on the features of the individual. In a high dimensional setting a non-parametric Kaplan-Meier estimator is typically used for 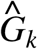, implicitly assuming the censoring distribution is independent from the observed features.

## Methods

### Retrieval and processing of RNA-Seq data

We employ RNA-Seq data from different tumor entities in the TCGA [26] provided in [27]. In total data for 24 cancer entities and a total of 9264 patients is available. We combine several different tumor entities into a total of three different risk groups in which we assume approximately the same baseline hazard: samples from glioblastoma multiforme (GBM) and brain lower grade glioma (LGG) are combined into one group, which we will refer to as GLIOMA. A second group includes tumor types related to pan-kidney diseases which are chromophobe renal cell carcinoma (KICH), kidney renal clear cell carcinoma (KIRC) and kidney renal papillary cell carcinoma (KIRP). We will refer to this group as KIPAN. The last group contains samples from breast invasive carcinoma (BRCA). In total, the resulting set includes 2555 individuals (1066 BRCA, 632 GLIOMA, 857 KIPAN), comprising the number of aligned reads per gene (gene-counts), survival time (time to event) and censoring indicator for every patient (see fig. 1 A). We removed patients where any of these values were missing. The whole dataset contains gene count information for more than 20.000 genes. To decrease the dimensionality of our data-set we employed a filter criterion not directly informative concerning survival, specifically the variance [28]. Our main results reported are achieved with the 3000 highest variable genes. These genes were additionally standardized by z-transformation before feeding them into the network.

### Training of deep neural networks

We trained multilayer feed-forward neural networks with 3000 input nodes, a varying number of hidden layers containing varying numbers of nodes per layer and a single output node (see fig. 1 B). In order to also capture the robustness of the investigated loss functions dependent on the network structure, we did not implement an extensive hyper-parameter optimization procedure in order to tweak our metrics to the best achievable score. Instead, we investigated 1000 randomly defined architectures. Specifically we first determined in each run a random network depth between 1 and 10 layers, and then generated a random width between 10 and 200 neurons for each layer (see fig. 1 B). The width of the layers were then sorted descending from input to output. On each of these generated network architectures a stratified 5-fold cross-validation was carried out, such that the ratio of samples from each group is reflected in the randomly selected training and testing subsets. On each of these generated architectures, we tested the stratified versus the non-stratified versions of the presented loss functions. Within each fold, all loss functions were trained and evaluated on the same training and testing subset of individuals. With a total of 1000 runs, this yields a total of 5000 training and testing phases for each loss function, with which we wanted to assure robustness for our results. We finished each training phase based on an early stopping criterion: training was stopped when the expected Brier score did not show any significant improvement on the test data anymore. As activation function, we used the rectified linear unit (ReLU), for parameter optimization we used ADAM [29]. No further regularization techniques were used. A link to our GitHub repository can be found in the supplementary material.

### Deep SHAP

One challenging part in using deep architectures is the interpretability of the fitted networks. In particular, contributions of individual feature are difficult to assess, and the approaches therefore often remain a black box. The SHAP approach was introduced as a unified way to address such issues [30]. It belongs to the class of additive feature attribution methods, which have an explanation model *g* that tries to explain the original model *f* through a linear function of binary variables:

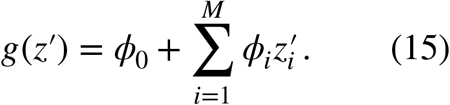

where *Z′* is a binary vector and *φ_i_* ∈ ℝ is the effect on the model output assigned to every feature. With results from cooperative game theory it can be shown, that there is only one possible explanation model to have the properties of local accuracy, missingness and consistency and solve the following equation:

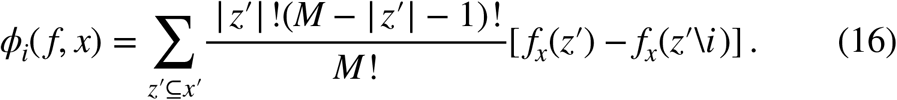

The solutions *φ_i_* are also known as Shapley values [31]. Adapting this proof to additive feature attribution methods yields the SHAP values with 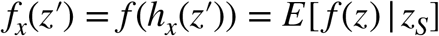
, and *S* the set of non-zero indexes in *Z′* Thus, SHAP values attribute to each feature the change in expected model prediction when conditioning on that feature [30]. When the employed model is non-linear or the features are not independent, the order in which features are added to the expectation matters, and SHAP values are obtained from averaging marginal contributions across all possible permutations. The exact calculation of these values can be computationally extremely expensive and therefore approximations are used. For the specific case of deep models [30] introduced *DeepSHAP*, which connect Shapley values with the DeepLIFT framework, which is a method for decomposing the output prediction of a neural network on a specific input by back-propagating the contributions of all neurons in the network to every feature of the input [32]. In our experiments, SHAP values are approximated through the DeepSHAP approach.

### Experiment – predicting survival from TCGA RNA-seq data

#### Classification and prediction performance dependent on loss function

We evaluated the stratified and non-stratified versions of the partial likelihood and the ranking loss based on 1000 different network architectures differing in the number of layers and nodes using 5-fold cross-validation. The prognostic performance was evaluated in terms of the prediction error and the C-index. For the C-index (fig. 2) median scores for all models range around 0.72. Employing stratified loss functions results in slightly larger, i.e. better C-indices. More striking are differences in the distribution of the C-indices for the different investigated loss functions. Using the ranking loss, stratified as well as non-stratified, results in more mass in the upper ranges. Comparing stratified and non-stratified loss functions shows, that stratified loss functions result in a smoother distribution across all runs. Especially when using the non-stratified ranking loss, three clear concentration points can be observed.

**Figure 2:**
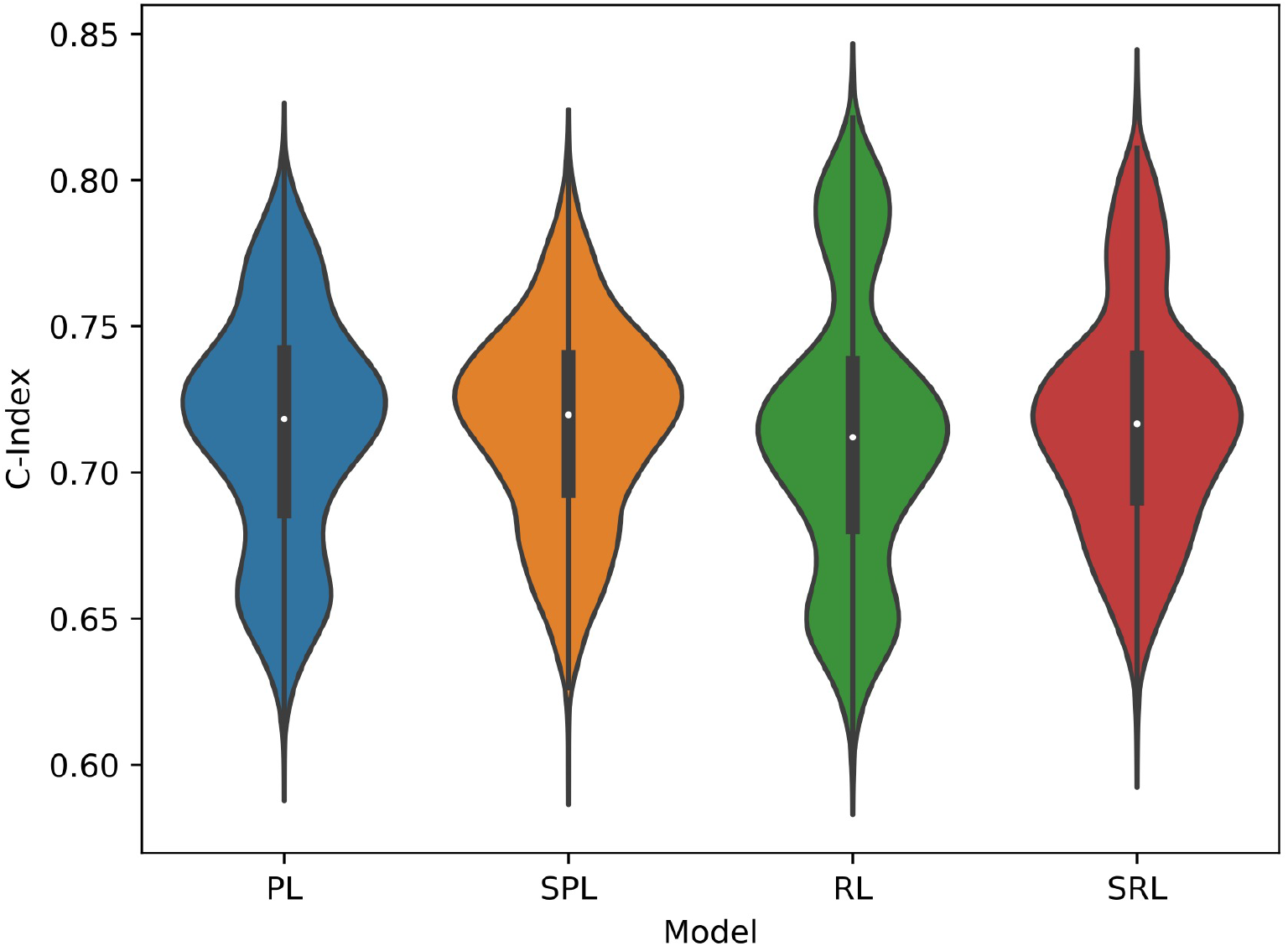
C-index dependent on loss function. 1000 different architectures were evaluated (See methods section for details). PL = partial likelihood, SPL = stratified partial likelihood, RL = ranking loss SRL = stratified ranking loss.

As already pointed out, the concordance index is not able to distinguish between well and badly calibrated models. Therefore we also evaluated the model performance in terms of the expected prediction error, also termed Brier score. Compared to the Kaplan-Meier estimator, chosen as performance baseline, almost all trained networks have a lower prediction error, irrespective of the employed loss function. Overall, we observe a strongly reduced prediction error, specifically integrated prediction error (iPEC) over time, when using the stratified loss functions (median iPEC 130.91 (PL) and 130.70 (RL) vs 125.79 (SPL) and 126.60 (SRL), Supplementary Table 1). When investigating the values of prediction error over time, we observe reduced error coinciding with increased observation time in networks trained based on stratified loss functions (fig. 3 – see e.g. the larger gap between the deep models and the Kaplan-Meier estimator observed for the stratified models, starting at day 500).

**Figure 3:**
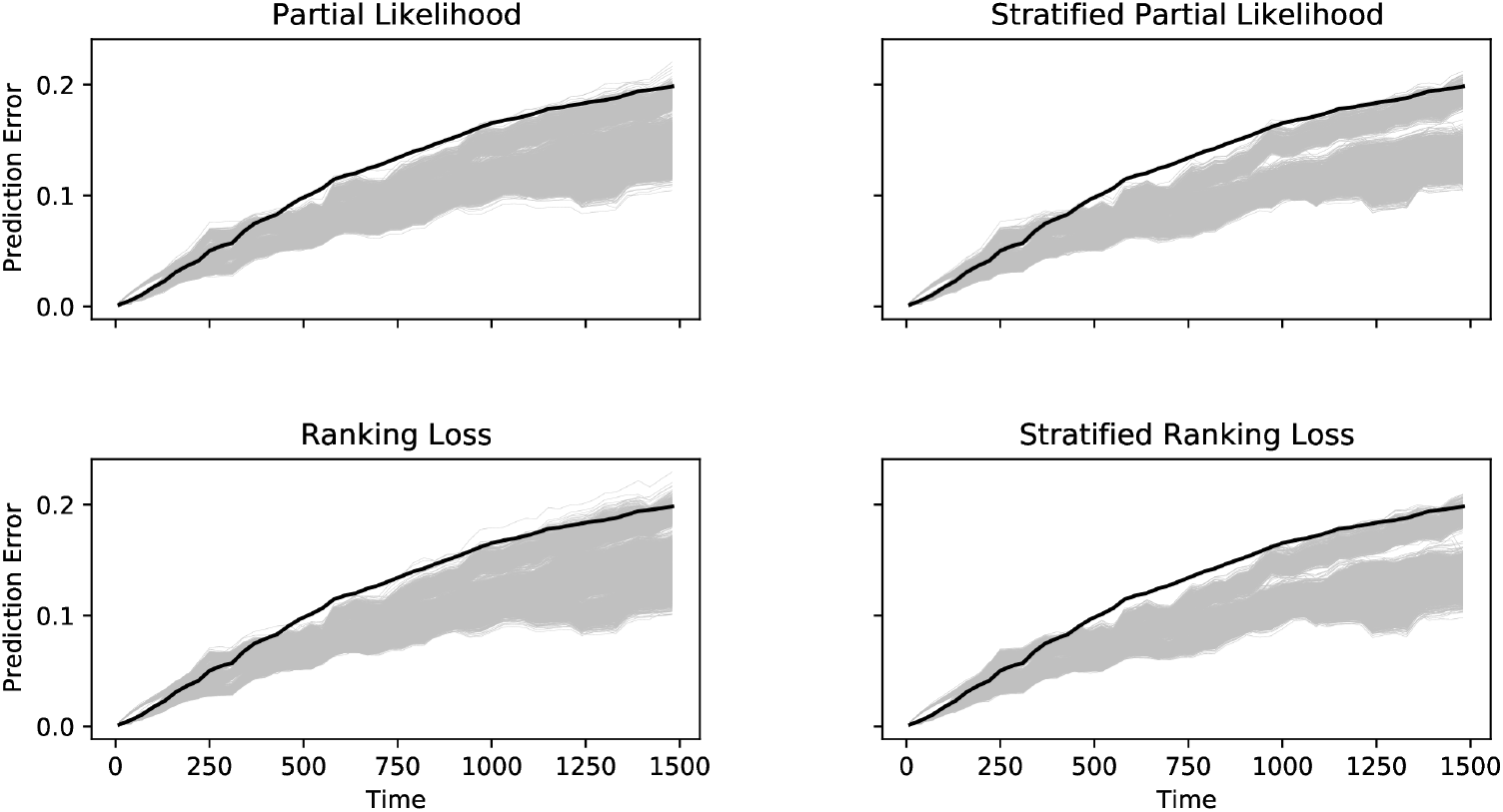
Prediction error curves (PEC) dependent on loss function. Prediction error is shown over time. The black line indicates the prediction performance of a non-parametric Kaplan-Meier estimation as a baseline. Each grey line corresponds to a model with a different architecture. 1000 different architectures were evaluated (See methods section for details).

#### Feature importance

To investigate whether the predictions made through the models are based on similar genes, we computed the marginal feature (gene) importance for each investigated patient. Specifically, we employ SHAP values. As the dependent variable is the predicted risk of experiencing the event, a positive SHAP value corresponds to a higher risk score. This means that genes with a higher density of SHAP values on the right hand side of the plots (fig. 4) can be considered as the high risk genes, as identified by the models. Based on the set of the genes with highest average SHAP values, calculated separately for all investigated loss functions, we investigate for each tumor entity whether the importance of genes is affected by the loss function. We observe mostly similar distributions of importance scores here (fig. 4 – BRCA, Supplementary Figure 1 – KIPAN, Supplementary Figure 2 – GLIOMA). Many of the genes are directly or indirectly associated with the prognosis of the different entities. For BRCA, e.g PAICS which is here associated with higher risk has been demonstrated to induce malignant proliferation of human breast cancer cells [33]. TCP1 has been demonstrated to be important for survival of breast cancer cells [34]. Similar results are obtained for KIPAN and GLIOMA. For instance, ALPK2 (Supplementary Figure 1 – KIPAN) promotes progression in renal cell carcinoma [35] and DLL3 (Supplementary Figure 2 – GLIOMA) is a therapeutic target to treat Glioma [36]. We also investigated the genes, where the average SHAP values were most strongly differentiated between the loss functions. Interestingly for BRCA, ADAM15, which is involved in prostate and breast cancer [37] received higher importance scores when employing the stratified loss functions (Supplementary Figure 3).

**Figure 4:**
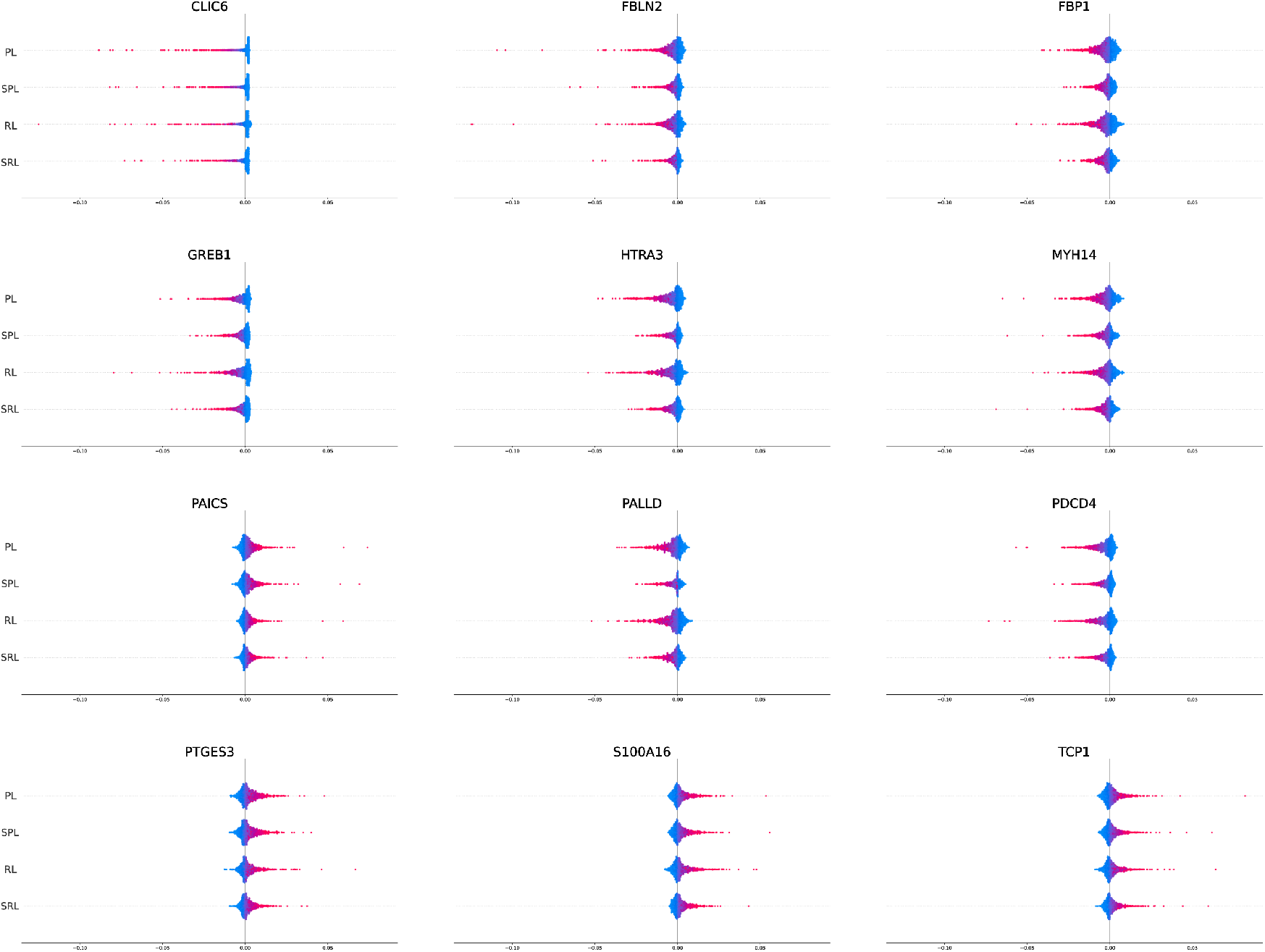
SHAP feature importance for BRCA patients. Shown is the union of the top five genes with highest average feature importance, extracted separately for each loss function. Each point corresponds to the marginal importance of the respective gene for predicting the relative survival of a single patient. The higher the absolute values, the more important the gene. Color relates to the expression strength of the gene, while magenta indicates higher expression and cyan indicates lower expression.

## Discussion and Conclusions

Here we investigated the performance impact of ignoring stratification when fitting models for time-to-event data building on deep neural networks, when different baseline hazard rates are expected in the data, and evaluated different approaches for incorporating stratification. Specifically, we investigated survival of cancer patients conditional on transcriptome profiles retrieved from tumor tissue. We also evaluated, whether the seemingly simple ranking loss, which is frequently employed with machine learning approaches, and the partial likelihood of a Cox proportional hazards model as a loss function benefit equally from a stratified variant. By investigating 1000 different network architectures, varying in the number of hidden layers and nodes per hidden layer, and by performing 5-fold cross validation, resulting in a total of 5000 runs, we assessed the robustness of the results.

We generally observed a higher C-index and lower prediction error in the stratified parameter estimation. However, the differences were more pronounced in terms of the prediction error. Especially with increased time, predictions benefit from the stratified procedure. These results are in concordance with previously known effects of violating the proportional hazards assumption [13]. Our findings also further demonstrate that the C-index as a evaluation criterion does not provide a comprehensive picture of model performance.

The loss functions used in our experiments maximize different objectives. While the estimates for the partial likelihood are directly related to survival probabilities, the parameters of the ranking losses represent an ordering function. Therefore, a loss function based on the partial likelihood should yield better results in predicting survival probabilities, while ranking losses should have an advantage in terms of the C-index, as they directly maximize the C-index. Yet, we obtained only slightly higher C-indices for the ranking loss, which is however not totally unexpected, since Cox proportional hazards model is known to approximately maximize the C-index [17].

While all loss functions resulted in rather similar importance scores for the genes that were most important for predicting relative survival, a gene with known importance in breast cancer (ADAM15), received higher importance when employing the stratified loss functions. This is a concrete example for avoiding bias through a stratified analysis.

One might argue that differences between the baseline hazards of groups should be more formally investigated. Yet, [38] showed that it is in fact very hard to show differences in baseline hazards given real, empirically derived data. Nevertheless, our results indicate, that employing stratified loss functions results in a substantial reduction in prediction error.

Furthermore, since we did not excessively tune hyper-parameters, our result might not reflect the full potential of the investigated models. However, given the multitude of different network architectures evaluated, the reported differences between stratified and non-stratified loss functions can be considered as robust.

Taken together, stratified analysis of observations that are suspected to originate from groups with different baseline hazards greatly enhances the prediction performance of models building on deep neural networks. This applies both to using the partial likelihood of Cox proportional hazards model or the ranking loss. Further, we show, that it is important to evaluate the predictive value of models for time-to-event data with an adequate measure, such as the Brier score.

### Key Points

1. The proportional hazards assumption allows for simple loss functions for training deep neural networks on time-to-event data for prognostic purposes. However, violating this assumption may lead to a biased estimation, which might negatively affect the prediction performance.
2. Here we investigate the benefit of using stratified loss functions for training deep neural networks on gene expression data with potentially different baseline hazards.
3. In an application on gene expression data from the TCGA, stratified loss functions result in better performance, specifically a slightly increased C-index and substantially lower prediction error.
4. While the marginal feature importance as revealed by deepSHAP suggests that most of the genes receive rather similar importance scores, there are examples for genes with known prognostic value that receive higher importance when employing stratified loss functions.

## Supplementary data

Supplementary data are available online at http://bib.oxfordjournals.org/. PyTorch code for stratified loss functions, Python scripts for recapitulation of the analyses and a Jupyter notebook which demonstrates the shown analyses are available on GitHub at https://github.com/fabriz-io/stratified_neural_network/.

## About the authors

Fabrizio Kuruc is a former bachelor student of the Institute of Medical Biometry and Statistics, Faculty of Medicine and Medical Center – University of Freiburg, Germany

Harald Binder is a professor and head of the Institute of Medical Biometry and Statistics, Faculty of Medicine and Medical Center – University of Freiburg, Germany

Moritz Hess is a postdoctoral associate at the Institute of Medical Biometry and Statistics, Faculty of Medicine and Medical Center – University of Freiburg, Germany

