## Supplementary data for "Stratified neural networks in a time-to-event setting"

### Stratified neural networks in a time-to-event setting - Supplementary Information

January 29, 2021

### Contents

### List of Figures

1 **SHAP feature importance for KIPAN patients.** Shown is the union of the top seven genes with highest average feature importance, extracted separately for each loss function. Each point corresponds to the marginal importance of the respective gene for predicting the relative survival of a single patient. The higher the absolute values, the more important the gene. Color relates to the expression strength of the gene, while magenta indicates higher expression and cyan indicates lower expression. . 2

2 **SHAP feature importance for GLIOMA patients.** Shown is the union of the top six genes with highest average feature importance, extracted separately for each loss function. Each point corresponds to the marginal importance of the respective gene for predicting the relative survival of a single patient. The higher the absolute values, the more important the gene. Color relates to the expression strength of the gene, while magenta indicates higher expression and cyan indicates lower expression. . 3

3 **SHAP feature importance for BRCA patients - difference.** Shown are the genes with the highest difference in average feature importance between the four investigated loss functions. Each point corresponds to the marginal importance of the respective gene for predicting the relative survival of a single patient. The higher the absolute values, the more important the gene. Color relates to the expression strength of the gene, while magenta indicates higher expression and cyan indicates lower expression. . 4

Supplementary Table 1: **Integrated prediction error curves for the four investigated loss functions.**

| Model | Median Integrated PEC | Mean Integrated PEC |
|---|---|---|
| Partial Likelihood | 130.91 | 132.96 |
| Stratified Partial Likelihood | 125.79 | 129.39 |
| Ranking Loss | 130.70 | 133.12 |
| Stratified Ranking Loss | 126.60 | 129.44 |

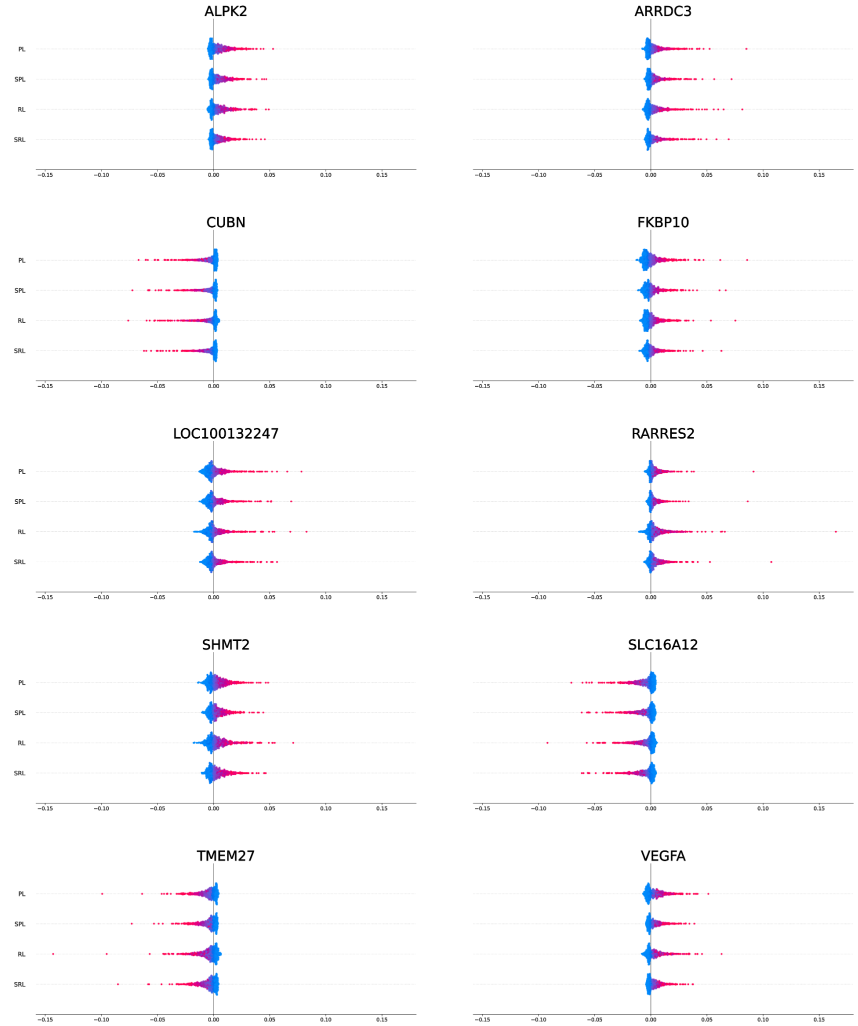


Supplementary Figure 1: **SHAP feature importance for KIPAN patients.** Shown is the union of the top seven genes with highest average feature importance, extracted separately for each loss function. Each point corresponds to the marginal importance of the respective gene for predicting the relative survival of a single patient. The higher the absolute values, the more important the gene. Color relates to the expression strength of the gene, while magenta indicates higher expression and cyan indicates lower expression.

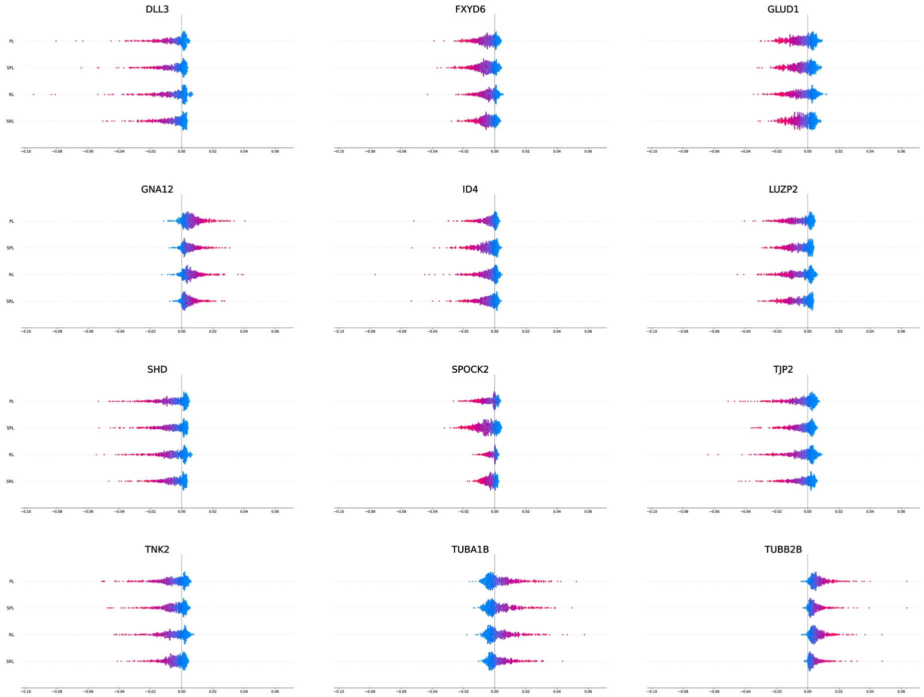


Supplementary Figure 2: **SHAP feature importance for GLIOMA patients.** Shown is the union of the top six genes with highest average feature importance, extracted separately for each loss function. Each point corresponds to the marginal importance of the respective gene for predicting the relative survival of a single patient. The higher the absolute values, the more important the gene. Color relates to the expression strength of the gene, while magenta indicates higher expression and cyan indicates lower expression.

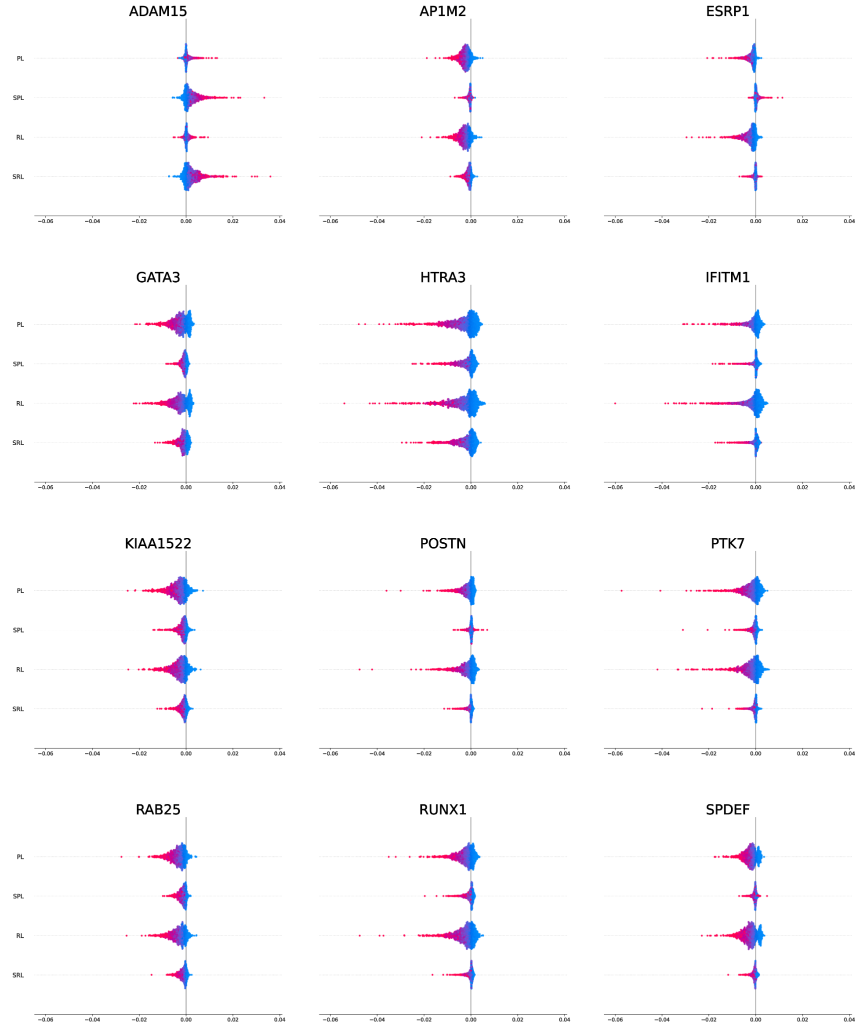


Supplementary Figure 3: **SHAP feature importance for BRCA patients - difference.** Shown are the genes with the highest difference in average feature importance between the four investigated loss functions. Each point corresponds to the marginal importance of the respective gene for predicting the relative survival of a single patient. The higher the absolute values, the more important the gene. Color relates to the expression strength of the gene, while magenta indicates higher expression and cyan indicates lower expression.
